# A new mRNA structure prediction based approach to identifying improved signal peptides for bone morphogenetic protein 2

**DOI:** 10.1101/2024.01.24.576995

**Authors:** Piers Wilkinson, Brian Jackson, Hazel Fermor, Robert Davies

**Author notes:** **Correspondence:** Piers Wilkinson.

## Abstract

**Background:** Signal peptide (SP) engineering has proven able to improve production of many proteins yet is a laborious process that still relies on trial and error. mRNA structure around the translational start site is important in translation initiation and has rarely been considered in this context, with recent improvements in *in silico* mRNA structure potentially rendering it a useful predictive tool for SP selection. Here we attempt to create a method to systematically screen candidate signal peptide sequences *in silico* based on both their nucleotide and amino acid sequences. Several recently released computational tools were used to predict signal peptide activity (SignalP), localization target (DeepLoc) and predicted mRNA structure (MXFold2). The method was tested with Bone Morphogenetic Protein 2 (BMP2), an osteogenic growth factor used clinically for bone regeneration. It was hoped more effective BMP2 SPs could improve BMP2-based gene therapies and reduce the cost of recombinant BMP2 production.

**Results:** Amino acid sequence analysis indicated 2,611 SPs from the TGF-β superfamily were predicted to function when attached to BMP2. mRNA structure prediction indicated structures at the translational start site were likely highly variable. The five sequences with the most accessible translational start sites, a codon optimized BMP2 SP variant and the well-established hIL2 SP sequence were taken forward to *in vitro* testing. The top five candidates showed non-significant improvements in BMP2 secretion in HEK293T cells. All showed reductions in secretion versus the native sequence in C2C12 cells, with several showing large and significant decreases. None of the tested sequences were able to increase alkaline phosphatase activity above background in C2C12s. The codon optimized control sequence and hIL2 SP showed reasonable activity in HEK293T but very poor activity in C2C12.

**Conclusions:** These results support the use of peptide sequence based *in silico* tools for basic predictions around signal peptide activity in a synthetic biology context. However, mRNA structure prediction requires improvement before it can produce reliable predictions for this application. The poor activity of the codon optimized BMP2 SP variant in C2C12 emphasizes the importance of codon choice, mRNA structure, and cellular context for SP activity.

## Background

Signal peptides (SPs), also known as signal sequences, are the region of secreted proteins that target them to a secretion pathway. In eukaryotes most secreted proteins use the signal recognition particle (SRP) secretion pathway, relying on short N-terminal signal peptides that induce co-translational translocation into the endoplasmic reticulum. The SRP pathway and its SPs have seen a great deal of basic research over several decades and much is understood of their function (reviewed elsewhere[1]), however SP sequences are highly diverse and it is not understood why some function better than others in specific contexts. Speculation that secretion may be a bottleneck in protein production was initially sparked by observations that in some contexts the amount of secreted protein did not increase proportionally with gene copy number, mRNA copy number or intracellular protein concentration[2, 3]. Subsequently many authors have successfully engineered SP variants or employed heterologous SPs to increase *in vitro* protein production for many proteins in a variety of hosts[4–13]. The approach has also seen application in the gene therapy and nucleic acid vaccine field, with various approaches proving successful *in vivo*[14–21].

Despite these successes, investigators must still rely on informed guesswork when selecting SP candidates. Often many need to be screened before an SP offering improved secretion is identified, and in some cases an improved SP is never identified[18, 22, 23]. A handful of SPs are known to be effective for multiple proteins and are commonly employed, though not always successfully. For example, the human serine protease 1 (trypsin) SP has been shown to improve secretion of the cytokines IL-25 and IFN-α2b[11, 12], yet did not improve secretion of the growth factor BMP2[18]. The human albumin SP has been shown to improve secretion of heavy and light chain immunoglobulins[5], but did not improve secretion of lysosomal enzymes NAGLU and GNS[24]. Perhaps an exception to this trend is the human IL2 (hIL2) SP, a common choice that has proved successful in various settings[14, 19–21, 25]. Currently we have been unable to find any studies where it failed to increase secretion, suggesting it is perhaps the most widely effective SP for *in vitro* eukaryotic expression currently known. However, it has not been universally employed and this may be a result of positive publication bias.

There have been some previous attempts to create systematic *in silico* approaches to eukaryotic SP selection or design, though they have seen little success and are not commonly used by the wider research community[26–28]. The Signal Peptide Secretion Efficiency Database is a promising alternative approach that attempts to address the issue by providing a repository of results for SP, gene and host combinations, though currently only contains data for a small number of proteins in prokaryotes[29]. Recent advances in *in vitro* screening methods show promise, though require large and complex screens that are not feasible for many researchers[30, 31]. There is still an unmet need for improved low-cost methods of choosing SP candidates.

A rarely considered aspect of SP behaviour is the influence of its nucleotide sequence on translation initiation. The SP nucleotide sequence lies at the 5’ end of coding sequence and contains the start codon and the last nucleotide of the Kozak sequence. Crucially, it is likely to form secondary structures with the latter segment of the 5’ UTR, the region containing the ribosomal attachment site. mRNA structure around the ribosomal attachment site is now established as an important mediator of translation initiation, with stable secondary structure in the region known to inhibit the process[32–36]. It therefore seems prudent to consider mRNA structure alongside amino acid sequence when selecting SP candidates. However, the large majority of SP optimization or substitution approaches do not do so, likely due to difficulties in predicting structure *in silico* or establishing it empirically. There have been a handful of attempts to integrate these factors in the past using *in silico* methods, though these were held back by difficulties in predicting mRNA structure and saw little success[37, 38]. The years since these attempts have seen rapid improvement in computational techniques due to advances in machine learning[39–43]. It was hoped the new generation of tools using these methods might offer improved performance for both amino acid sequence based SP prediction and mRNA structure prediction.

Bone Morphogenetic Protein 2 (BMP2) is an osteogenic growth factor in the TGF-β family that plays important roles in bone development, homeostasis and healing[44–49]. BMP2 has seen extensive use in regenerative approaches for bone healing and other orthopaedic applications, both as a recombinant protein and a gene therapy[50, 51]. Recombinant human BMP2 (rhBMP2) has been used in the clinic for spinal fusion and fracture treatment for 20 years, though issues have emerged over the high incidence of side effects due to the initial high dose required to counteract the protein’s poor biological half-life[52, 53]. Gene therapies hope to combat this issue by inducing controlled medium-term production of the protein in the defect itself. Despite decades of research no gene therapies for bone regeneration are currently approved for clinical use, largely due to safety concerns around viral vectors and the poor efficacy of non-viral alternatives[54]. It is hoped that methods to improve the effectiveness of non-viral vectors such as SP engineering have the potential to enable clinical translation of non-viral BMP2 gene therapies. Only one prior publication has attempted to identify improved SPs for BMP2 using a non-systematic approach, failing to identify any SPs offering improved secretion[18].

Here we present a new method to identify SPs to improve BMP2 secretion. The method employs various up-to-date computational tools from other authors to predict SP activity based on amino acid and nucleotide sequences, with particularly attention paid to the predicted mRNA structure at the ribosomal attachment site. A subset of the top results and two manually selected sequences of interest were then tested *in vitro* in two cells lines, with protein secretion and osteogenic activity investigated. We report the strengths and weaknesses of the technique and suggest several approaches to improve the method.

## Methods

### Acquisition of signal peptide amino acid sequences and creation of the *in silico* fusion peptide library

The protein sequences for all identified members of the TGF-β superfamily in all species were downloaded from the UniProt database as an XML file on 13/10/2022 (URL: https://www.uniprot.org/uniprotkb/?query=family:%22TGF-beta+family%22&sort=score)[55]. The SP amino acid sequences of proteins annotated with SPs were isolated and attached to the hBMP2 propeptide amino acid sequence, creating an *in silico* library of fusion proteins. Nucleotide sequences corresponding to the SP amino acid sequences were retrieved by isolating source database information and accession numbers from the UniProt data. The various nucleotide sequence databases cited (NCBI nucleotide, EBI-ENA, Ensembl, WormBase ParaSite[56–59]) were then accessed by their application programming interfaces to retrieve the relevant sequence data. After retrieval, the nucleotide sequence dataset was subjected to an availability and validity check using the following criteria: nucleotide sequence successfully retrieved, sequence in frame, sequence begins with a start codon, and translated nucleotide sequence matches the UniProt protein sequence.

### Prediction of signal peptide function and localisation with SignalP and DeepLoc

The *in silico* fusion proteins were analysed with signal peptide prediction software SignalP 6.0 (Technical University of Denmark) to predict if they would be recognised by secretion machinery[60]. The fast model was chosen due to substantially reduced completion time for the large data set used here. SignalP gives a score for the predicted likelihood of the sequence containing a functional eukaryotic SRP pathway signal peptide (called the Sec/SPI score) between 0 and 1, with sequences given higher scores considered more likely to contain an SRP signal peptide. An exclusion threshold of Sec/SPI < 0.5 was set.

The signal peptide targeting predictor DeepLoc 2.0 (Technical University of Denmark) was used to predict the localisation targets of the SPs[61]. DeepLoc provides predictions of localisation to various intracellular targets and the extracellular space, giving a score of 0 to 1 for each location with higher values indicating increased confidence in the prediction. An exclusion threshold of a <0.5 for extracellular localisation was set.

### Manually selected sequences

The expression optimisation tool “Translation Initiation coding region designer” (TISIGNER, https://tisigner.com/tisigner) was used to design a codon optimised version of the endogenous hBMP2 SP. TISIGNER prioritises minimal mRNA 5’ opening energy rather than using a more standard tRNA availability based approach (see Bhandari *et al.* for a full description of the method[62]). Host organism was specified as “Other” and the following promoter and 5’ UTR sequence from the intended pVax plasmid vector was entered to allow mRNA structure prediction (5’ – 3’ orientation):

GTGATGCGGTTTTGGCAGTACATCAATGGGCGTGGATAGCGGTTTGACTCACGGGGATTTCCAAGTCTCCACCCCATTGACGTCAATGGGAGTTTGTTTTGGCACCAAAATCAACGGGACTTTCCAAAATGTCGTAACAACTCCGCCCCATTGACGCAAATGGGCGGTAGGCGTGTACGGTGGGAGGTCTATATAAGCAGAGCTCTCTGGCTAACTAGAGAACCCACTGCTTACTGGCTTATCGAAATTAATACGACTCACTATAGGGAGACCCAAGCTGGCTAGCGTTTAAACTTAAGCTTGGTACCGAGCTCGGATCCACTAGTCCAGTGTGGTGGAATTCGGCTTGCCACC

The full hBMP2 mature transcript nucleotide sequence from the NCBI nucleotide database (Accession: NM_001200.4) was then entered for optimisation.

The signal peptide nucleotide sequence from the hIL2 mRNA was taken from the hIL2 references sequence on NCBI nucleotide (Accession: NM_000586.4).

### Creation of *in silico* predicted mRNAs and structure, stability and opening energy prediction

To predict the structure and stability of the fusion proteins’ mRNAs when expressed from the intended pVax_BMP2 based plasmid vector (sequence available at https://www.ncbi.nlm.nih.gov/nuccore/MK433563.1), *in silico* predicted mRNAs were created. In 5’ to 3’ orientation these contained the 5’ UTR of pVax_BMP2, the SP of interest, the hBMP2 propeptide, the pVax_BMP2 3’ UTR, the BGH poly-A sequence to 20bp downstream of the AATAA polyadenylation site, then a 150bp poly A tail. The resulting predicted mRNA sequences were ∼1.7kb in length.

The RNA secondary structure predictor MXFold2 (Department of Biosciences and Informatics, Keio University, Japan) was used to predict the secondary structures and calculate the stability of the predicted mRNAs[63]. Stability was quantified with minimum free energy (MFE) values, which account for the predicted energetic contribution of every feature in the predicted secondary structure[63]. To calculate opening energy the MXFold2-generated secondary structures were submitted to RNAeval (see ViennaRNA package[64]) to provide the predicted energetic contribution of each bond in the MXFold2 generated structure. Opening energy was predicted by extracting the energy values for the ±15nt segment surrounding the start codon (previously established as a relevant window by others[33–35]) using a python script. A simple metric was devised to simultaneously consider the predicted opening energy and total mRNA stability values, generating a metric termed SP score.

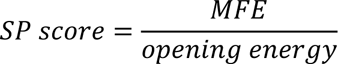

Higher SP scores indicate larger predicted MFE and smaller predicted opening energy; the hypothetically favourable combination of properties for maximising expression.

### Additional *in silico* analysis

RNA structure visualisation was performed with the Forna web app. (http://rna.tbi.univie.ac.at/forna/). See Kerpedjiev et al. for details[65]. Nucleotide and amino acid sequence alignments were performed in SnapGene (SnapGene software, www.snapgene.com). The final 7 candidates and the endogenous hBMP2 SP were submitted to the SignalP 6 web app (https://services.healthtech.dtu.dk/services/SignalP-6.0/) to provide predictions of region boundaries[60]. “Organism: Eukarya”, “Output format: Long” and “Model mode: Slow” were specified prior to running the model.

### Plasmids

The plasmids pVax_BMP2 and pVax_BMP7 were kindly provided by Dr Georg Feichtinger (see Feichtinger *et al.* for details[66]). The pVax_GFP control plasmid was produced from pVax_BMP2 by restriction cloning. Briefly, the BMP2 CDS was removed and replaced with EGFP excised from the pCAG_GFP plasmid previously purchased from Addgene (https://www.addgene.org/11150/). Novel SP fragment synthesis, restriction cloning into pVax_BMP2 and endotoxin free maxi-prep scale up were outsourced using the Genewiz TurboGENE 7 Day service (Azenta life sciences).

### Cell culture

HEK293T cells were kindly provided by Dr Brian Jackson. The cells were maintained in DMEM high glucose (D6429, Sigma), 10% FBS (EU-000-F, Seralab), 2 mM L-Glutamine (G7513, Sigma), 1% v/v penicillin/streptomycin. C2C12 cells were purchased from ATCC (CRL-1772). The cells were maintained in DMEM high glucose with 4mM L-Glutamine and 5% FBS, without antibiotics. Both lines were maintained at 37^0^C in a 5% CO2 atmosphere and passaged twice a week during maintenance periods.

HEK293T cells were seeded in 24-well plates at 3x10^5^ cells/well in 1ml of complete DMEM, 24 hours prior to transfection. Cells were transfected using Lipofectamine 3000 (L3000001, Thermo Fisher Scientific) at the manufacturer recommended high concentration. After 24 hours an additional 1ml of complete DMEM was added to each well to ensure the media was not exhausted before harvesting. Media was harvested 48 hours after transfection. SP panel experiments contained 9 groups with each transfected with one of the following plasmids: the top 5 results from the in silico screen, the two manually selected SP sequences (SP hBMP2 TISIGNER and SP hIL2), a pVax_BMP2 positive control and a pVax_GFP negative control. Transfections were always performed in technical duplicate with two wells per group.

C2C12 cells were seeded in 24 well plates at 6x10^4^ cells/well in 1ml of complete C2C12 media. The cells were transfected after 24 hours as described above. Experimental groups were the same as those in the HEK293T experiment with the addition of a co-transfection pVax_BMP2 +pVax_BMP7 osteogenesis positive control group (effectiveness previously established by others[66]). In this group 250ng of each plasmid was used per well to match the 500ng used in the other groups. 24 hours after lipofection the cells were media changed with 1ml of complete media supplemented with 10µg/ml heparin (H3149, Sigma Aldrich) to reduce interaction with heparan sulphate proteoglycans that would otherwise rapidly clear hBMP2 from solution[67, 68]. After a further 24 hours (48 hours after lipofection) the media was harvested for BMP2 ELISA and new heparin-supplemented media added. The plates were maintained in heparin-supplemented media until 7 days post transfection, with media changes every 2/3 days. On day 7 the wells were washed with PBS and the plates stored at - 80^0^C for up to 48 hours prior to quantitative alkaline phosphatase (ALP) assay.

### hBMP2 ELISA

hBMP2 capture and biotin-conjugated detection antibodies, CHO-derived rhBMP2 standard and streptavidin-HRP working solution from the hBMP2 DuoSet ELISA system (DY355, R&D systems) were used for all experiments. Nunc MaxiSorp ELISA plates (M9410, Merck), 10% BSA ELISA reagent diluent/blocking solution concentrate (DY995, R&D systems) and TMB substrate (421501, Biolegend) were purchased separately and used for all experiments. ELISAs were performed according to the hBMP2 DuoSet ELISA system manufacturer recommendations. rhBMP2 standard curves were made up in the appropriate complete media. A Wellwash microplate washer (5165000, Thermo Fisher Scientific) was used for all washes.

### Quantitative ALP assay

Nitrophenol phosphate (NPP) quantitative ALP assays were performed to assess osteogenesis in the C2C12s seven days after transfection. Cells were washed with PBS then lysed with 100ul lysis solution (0.5% Triton X-100 in deionized water) per well with 250RPM radial shaking for 1 hour. 10ul from each well was transferred to a new 96 well plate, then a 90ul of NPP working solution (5mM NPP (4876, EMD Millipore), 0.5M 2-AMP (A9199, Sigma Aldrich), 2mM MgCl2 (25108.260, VWR chemical), pH 10.3 in deionised water) was added to each well. A 4-Nitrophenol (4NP) end-product standard curve of 10-200µM (4NP to concentration(1048, Sigma Aldrich), 0.5M 2-AMP, 2mM MgCl_2_, pH 10.3 buffer) was run simultaneously to allow calculation of the final 4NP concentration in each well. Plates were incubated at room temperature for 45 minutes then absorbances at 405nm and a 600nm wavelength control were measured using a microplate spectrophotometer (Multiskan GO microplate reader, Thermo Fisher).

### Statistics

ELISA and ALP data were normalised to the positive control group prior to analysis. For ELISAs the positive control was the pVax_BMP2 group, while for ALP assays the positive control was the pVax_BMP2+pVax_BMP7 group. The normalised data were then tested for normality using Shapiro-Wilk tests. All data were found to be normally distributed, and significance was tested using one-way ANOVAs with selected Dunnett’s multiple comparison tests between the positive control and other groups to improve power. All statistical tests were performed in GraphPad Prism 9.3.1 (GraphPad Software).

## Results

### Results from the *in silico* pipeline

The initial TGF-β family dataset retrieved from UniProt contained 34,861 records. 19,250 protein sequences were found to be annotated with a confirmed or likely signal peptide and were taken forward. Nucleotide sequence retrieval and quality control checks (sequence in frame, sequence begins with a start codon, translated nucleotide sequence matches the UniProt protein sequence) were then performed. 1,344 sequences failed one of these criteria and were removed, leaving 17,906 to be taken forward. An SP nucleotide sequence duplicate removal step was then performed, with 7,545 sequences found to be duplicates and removed. 10,361 unique and matched nucleotide/amino acid sequence pairs remained and were used to create the *in silico* fusion protein and predicted mRNA sequence libraries used for further analysis. The *in silico* fusion protein library was subjected to SP activity and localisation prediction. SignalP analysis indicated that 3,203 of the fusion proteins were predicted to have a low chance of secretion and were excluded, leaving 7,158 sequences remaining. DeepLoc analysis indicated all the fusion proteins were predicted to show extracellular secretion, therefore none were excluded. Nucleotide sequences for these fusion proteins were then analysed for weak Kozak sequences. 4,547 sequences were found to have non-optimal Kozak sequences and were excluded. This left a final set of 2,611 sequences for further analysis. See figure 1 for an overview of the filtering process.

**Figure 1:**
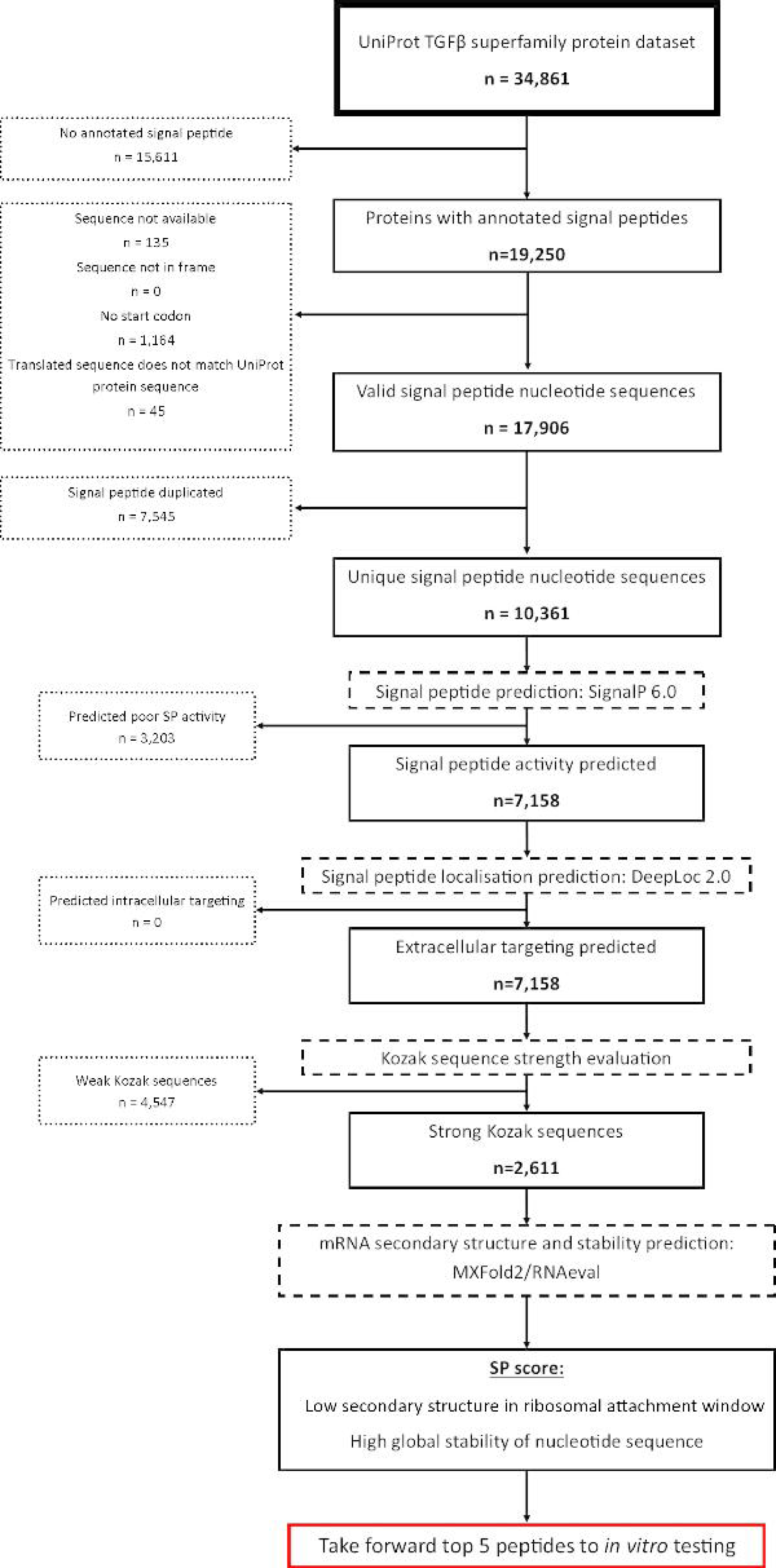
Overview of the filtering process employed in the *in silico* pipeline. Initially ∼35,000 TGF-β superfamily sequences were retrieved, though only ∼19,000 were predicted to contain SPs. Of these, ∼10,000 were found to be valid records and to have unique nucleotide sequences. Existing computational methods indicated ∼7,000 were predicted to still function and to target the extracellular space when attached to BMP2. ∼2,500 were found to have strong Kozak sequences and were taken forward to mRNA structure prediction. Sequences with the least structure at the translational start site but high global stability were thought to be preferable. The top 5 sequences from the pipeline and two manually selected alternatives were taken forward to *in vitro* work.

The secondary structures of the predicted mRNA sequences for the remaining SP-fusions were predicted with MXFold2. Predicted opening energies varied substantially across the final data set (mean: -343.3 kcal/mol, SD: 106.6), while MFE was much less variable (mean: -451.9 kcal/mol, SD: 11.2). The most promising 5 results from the in silico screen according to their calculated SP score are displayed in table 1. The sequences were all derived from different vertebrate species. Three sequences were derived from ARTN or closely related unknown protein orthologs, in all cases from mammalian species. The two additional identified proteins were BMP15 and GDF-10 orthologs, from an avian and piscine species respectively. There was modest variation in SP length, ranging from 22 to 44 residues in length (note the native hBMP2 SP is 23 residues long). The MFEs of the 5 sequences showed little variation (mean: -466.7 kcal/mol, SD: 19.6). Opening energies were more variable but covered only a small fraction of the range seen across the whole dataset (mean: -72.6 kcal/mol, SD: 13.6). Sequence alignments revealed that there was limited similarity between sequences from distantly related species and proteins. There was unsurprisingly strong sequence similarity between the 3 artemin orthologs (see figure 2A and 3A), with all displaying a ∼25bp insertion. All three sequences were predicted to show the same mRNA secondary structure in the ribosomal binding site window (see figure 2B). A loose leucin-rich motif was conserved in all sequences (see figure 3A).

**Figure 2:**
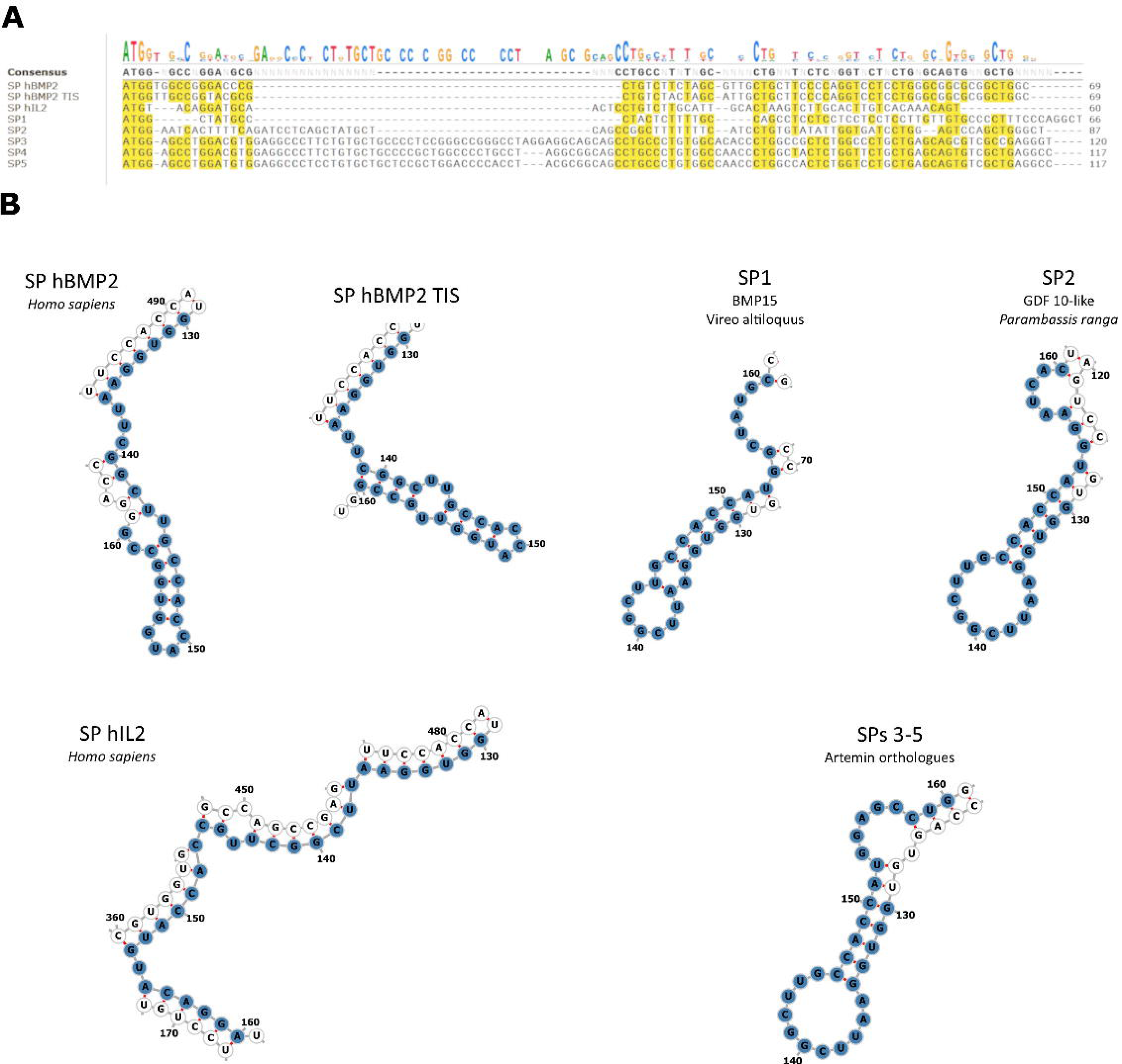
Further nucleotide sequence analysis A) Nucleotide sequence alignment of the final 7 SP sequences. Note the moderate variety with few positions being strongly conserved across the final set. A ∼25bp insertion can be seen in the SPs from Artemin related genes (SPs 3-5), making them noticeably longer than the other sequences. B) Predicted mRNA secondary structures at the ribosomal attachment site. The ±15bp window is highlighted in blue, with start codon found at positions 151-153.

**Figure 3:**
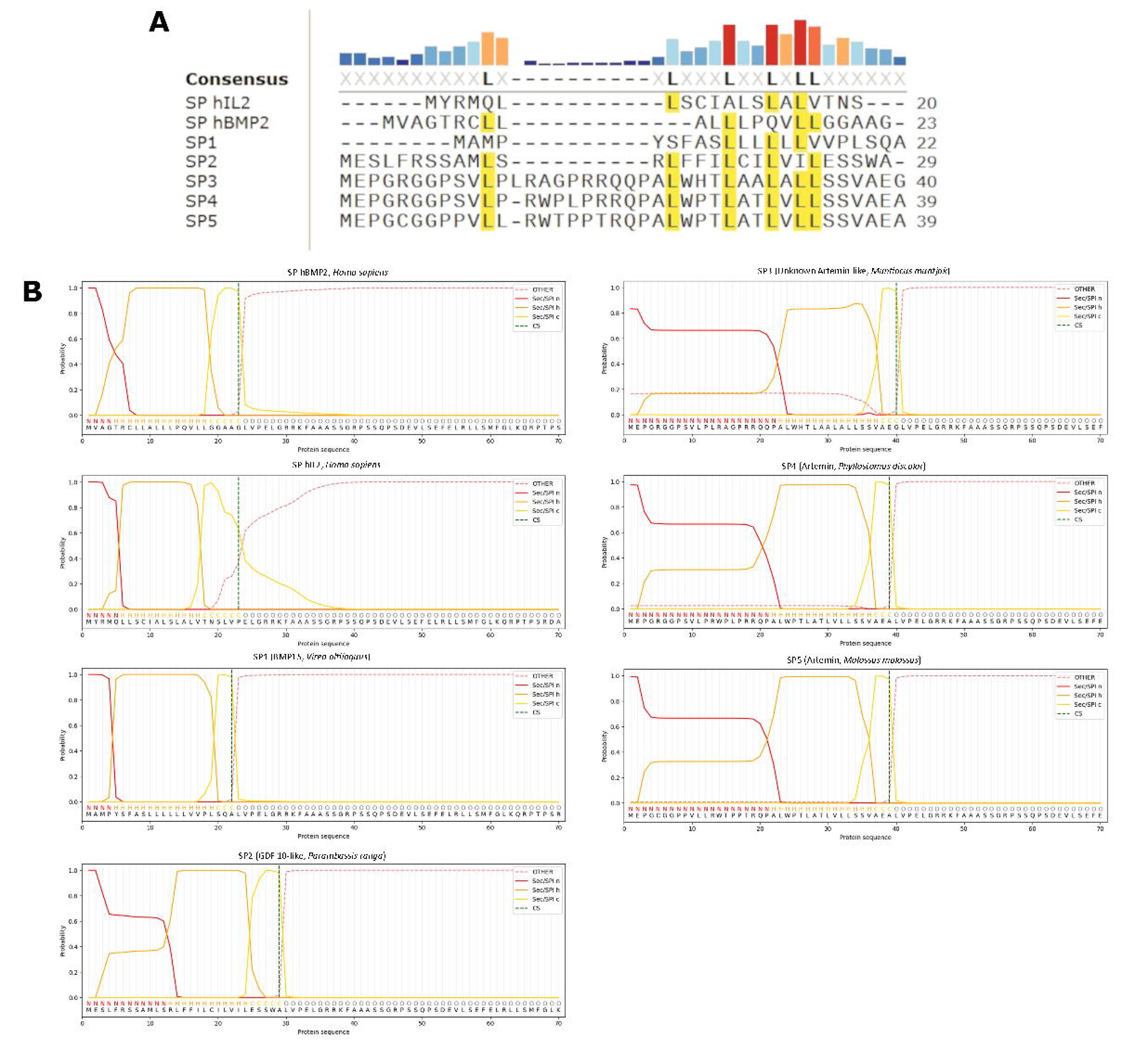
Further protein sequence analysis A) Protein sequence alignment of the final 7 SP sequence. Note the series of conserved leucine residues, the only conserved element across the final panel. Unsurprisingly the three SPs from Artemin related genes (SPs 3-5) were highly similar. B) Detailed SP region boundary predictions from the SignalP long model. The Artermin related SPs displayed elongated N-regions in comparison to the rest of the set, corresponding to the position of the insertion in the nucleotide sequences (see figure 2A). Also note the leucine motif from the alignment is contained within the SP H-region, thought to be important for maintaining hydrophobicity and ensuring the alpha helical conformation required for transmembrane function.

**Table 1:**
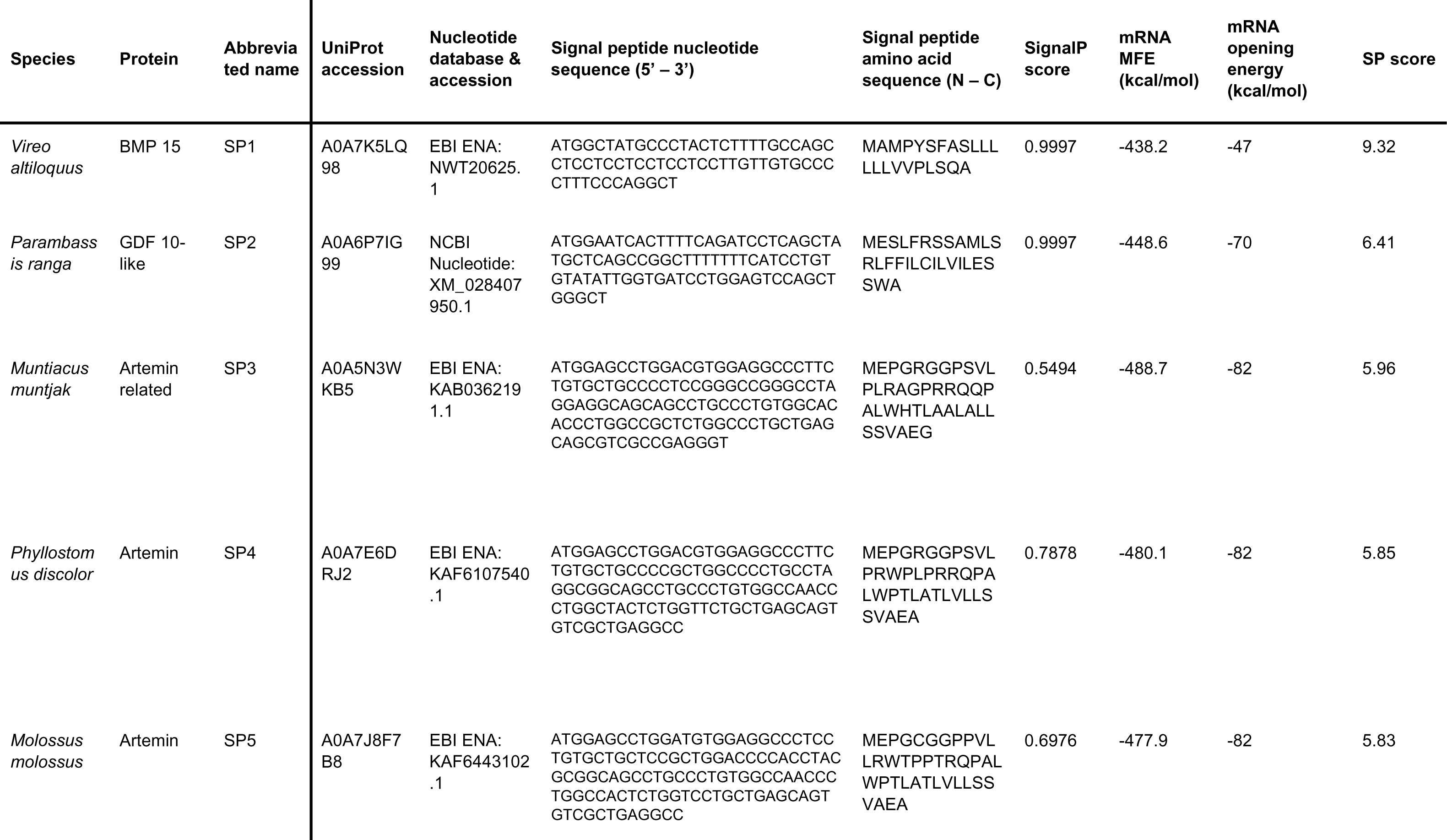
Overview of species, protein, sequence data and calculated RNA properties for the top results from the *in silico* screen.

The endogenous hBMP2 SP and the two manually selected sequence, SP hBMP2-TIS and SP hIL2, showed similar MFEs but large opening energies compared to the *in silico* pipeline top 5, with IL2 being particularly high (see table 2) due to predicted low self-complementarity (see figure 2B). Interestingly the hBMP2-TISIGNER sequence, supposedly codon optimised for lowered opening energy, was predicted to have a marginally increased opening energy when compared to the native sequence. This likely results from differences in the methods employed by TISIGNER and MXFold2/RNAEval when predicting structure and assigning bond energy values. All three sequences showed low SP scores, resulting in them acting as useful counterpoints to the top in silico candidates when testing the usefulness of the metric in vitro.

**Table 2:**
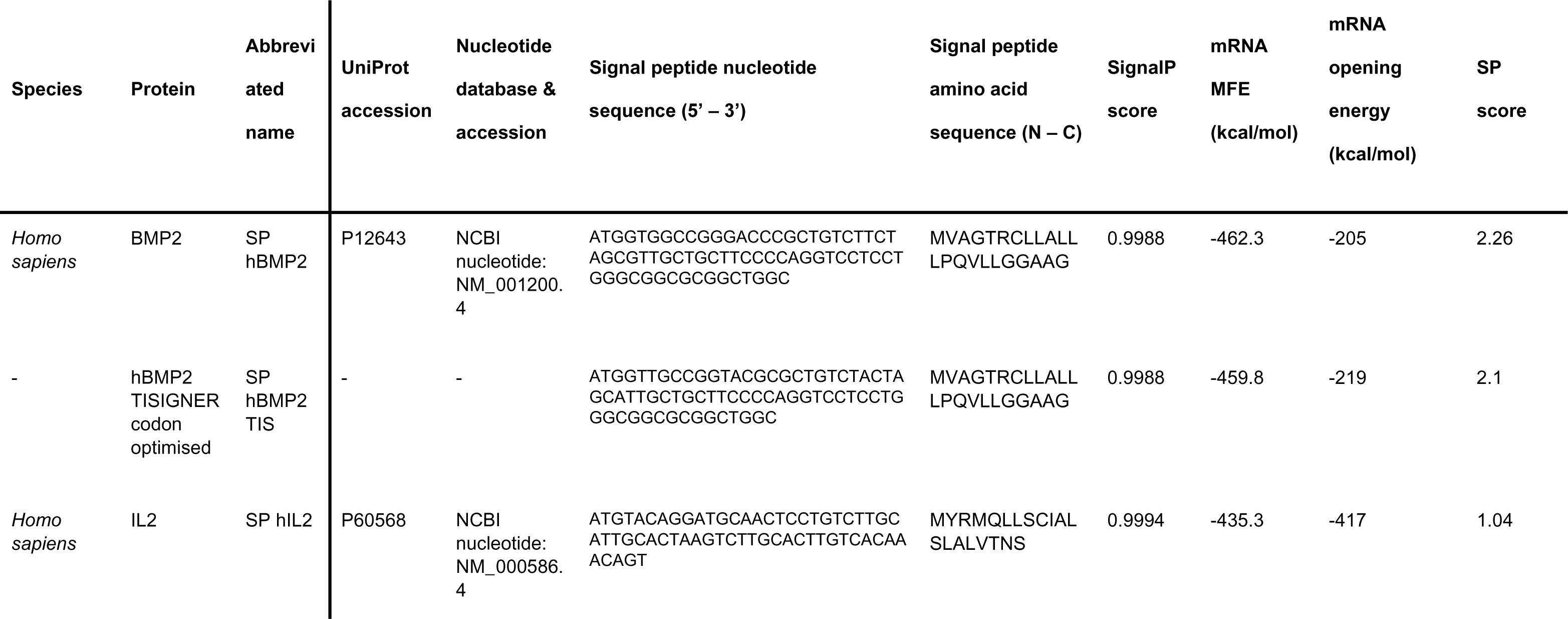
Overview of species, protein, sequence data and calculated RNA properties for the manually selected sequences.

The final seven candidates to be taken forward to in vitro testing were subjected to additional *in silico* analysis. The amino acid sequences were re-submitted to SignalP using the slow model for an analysis of predicted SP region boundaries. The four sequences that were noticeably longer than the native hBMP2 SP (GFD-10 from *P. ranga* & the artemin orthologs) were predicted to have much longer N-regions than the other sequences (see figure 3B). The N-region of hBMP2 is only four residues long, while that of *P. ranga* GDF-10 was 12 residues and the artemin orthologs were 21-22 residues. Additionally, the SP cleavage site for the hIl2 SP was predicted with less confidence than the other sequences and was 3 residues downstream of the true SP/preprotein sequence boundary (see figure 3B). The leucine motif identified in the alignment was predicted to be part of the H-region(see figure 3B), an established loose feature of mammalian SPs[1].

### *In vitro* validation in two cell lines

ELISA results from HEK293T conditioned media showed that the signal peptides identified by the *in silico* work and the manually selected sequences of interest did not induce a significant increase in hBMP2 secretion versus the pVax_BMP2 positive control (see figure 4A). The *in silico* selected signal peptides all showed small but non-significant increases versus the pVax_BMP2 positive control, while the manually selected SPs showed small and non-significant decreases. Data were highly variable across all groups, with the exception of the pVax_GFP negative control. There appeared to be a minor blanking issue as small negative hBMP2 concentrations were measured for the negative control across all replicates. This was thought to be a minor issue as these values were only a small fraction of that detected in the other samples. It was suspected this was caused by slight differences in blocking behaviour between the fresh complete media used for the standard curve and the conditioned media that made up the samples. C2C12s conditioned media ELISA results showed significant decreases versus the pVax_BMP2 positive control for SPs 1 & 2, SP hBMP2 TISIGNER and SP hIL2, while in silico SPs 3,4 & 5 showed non-significant decreases (see figure 4B). After 7 days ALP activity was again at background levels in all groups except the osteogenesis positive control (see figure 4C).

**Figure 4:**
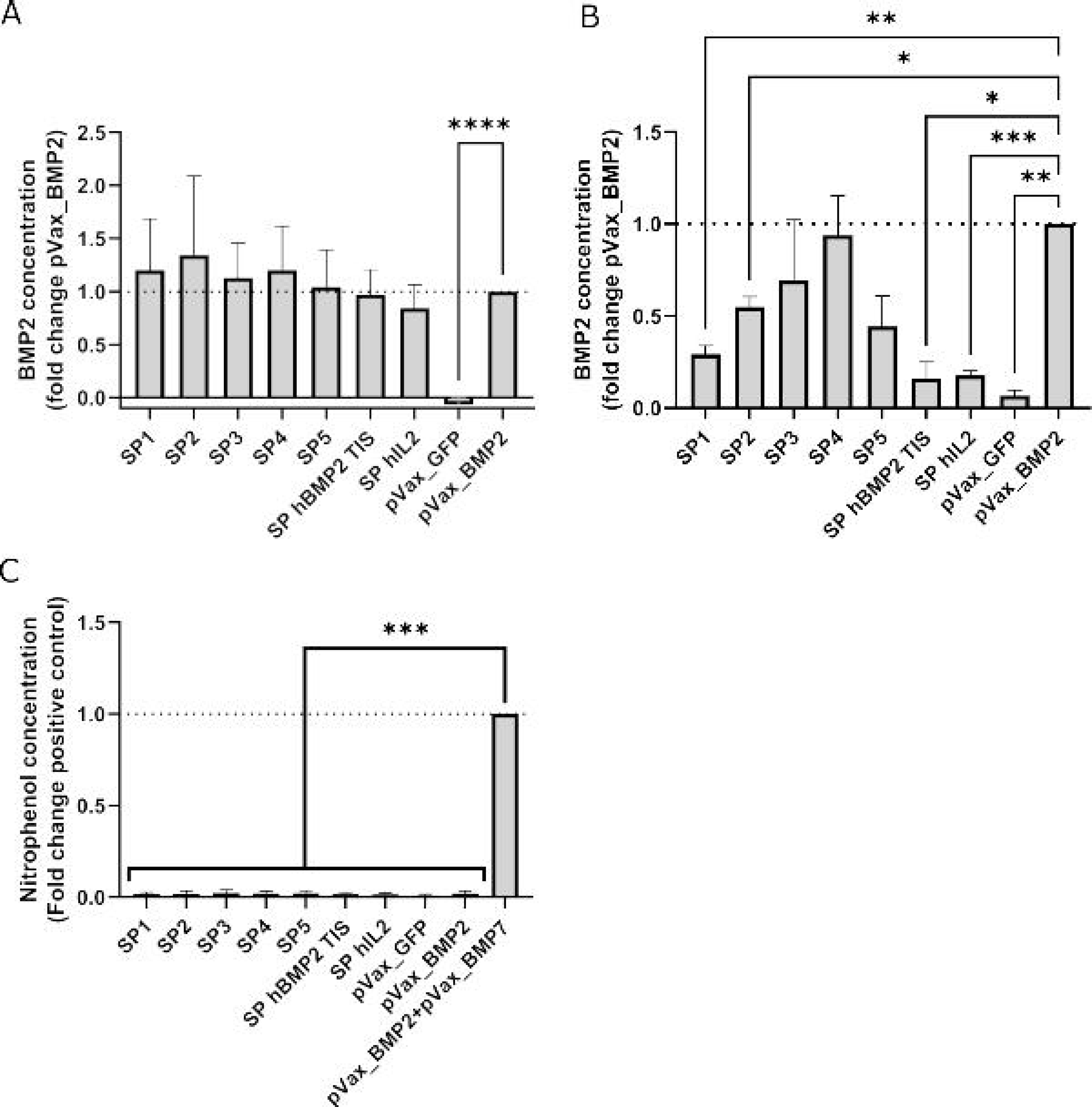
*In vitro* validation data A) hBMP2 ELISA data from HEK293T cells. All computationally selected sequences showed non-significant increases in secretion. N=7±SD, **** = p<0.0001 B) hBMP2 ELISA data from C2C12 cells. All computationally selected sequences showed decreased secretion versus the positive control, with several showing significant decreases. Particularly notable were the sequences that showed major decreases in the C2C12 results compared to the HEK293T experiment. These were SP1, SP2, SP hIL2 and SP hBMP2 TIS. N=3±SD. * = p<0.05, ** = p<0.01, *** = p<0.001 C) ALP assay data from C2C12 cells. All groups except the positive control showed background levels of ALP activity, indicating the novel SPs were not sufficient to induce osteogenesis in this context. N=3±SD, *** = p<0.001

Regression analysis of SP scores and ELISA data indicated that there was a significant correlation between SP score and BMP2 secretion in HEK293T (see figure 5A; gradient = 0.0509, R^2^ = 0.096, F(1, 47) = 4.976, p = 0.0305). No correlation was found between SP score and the heparin supplemented C2C12 ELISA data (see figure 5B).

**Figure 5:**
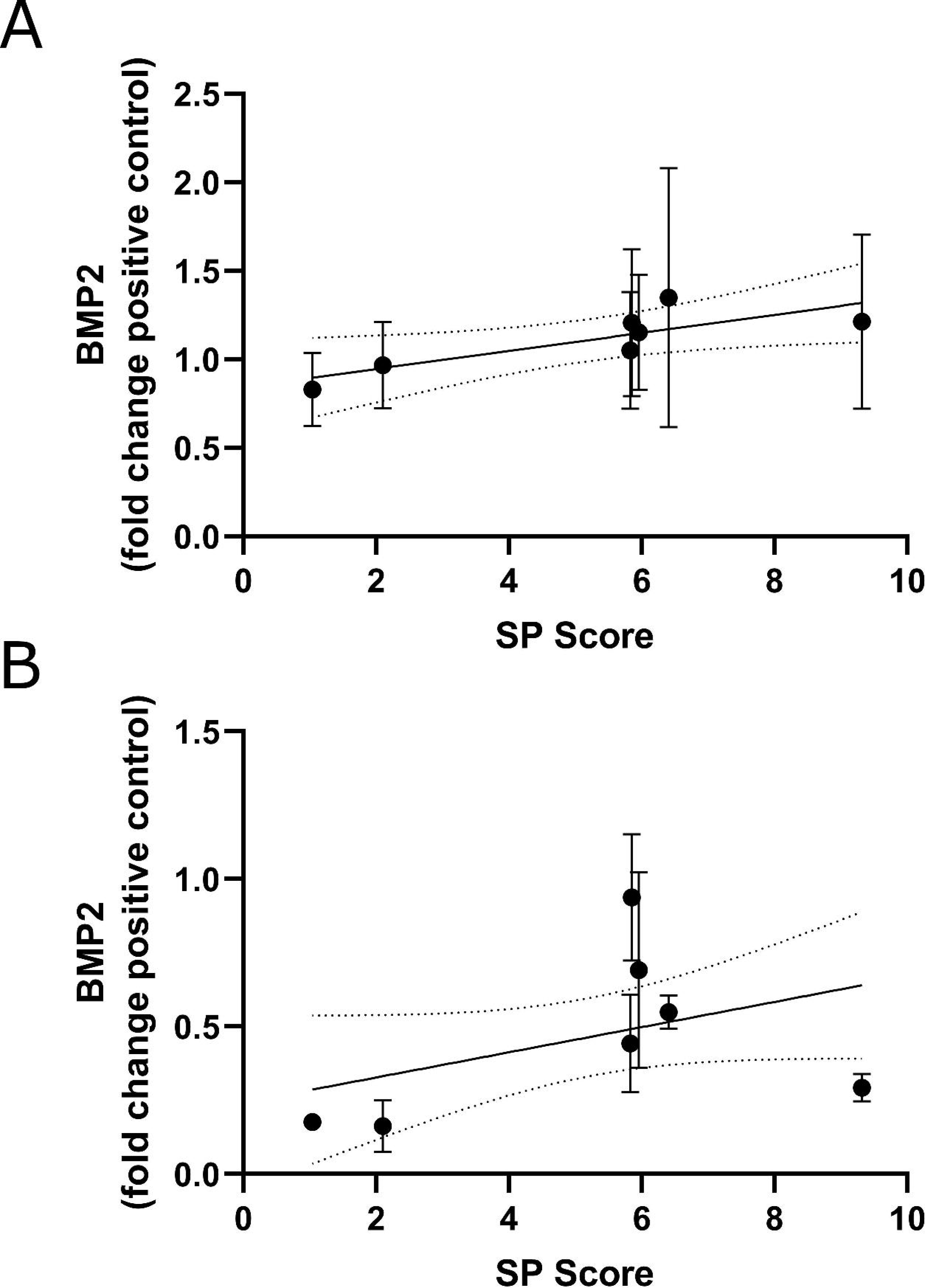
Linear regression analyses of the SP score values and ELISA data A) ELISA data from the HEK239T experiment. Correlation was found to be significant (gradient = 0.0509, R2 = 0.096, F(1, 47) = 4.976, p = 0.0305) B) ELISA data from the C2C12 experiment. No significant correlation was found (gradient = 0.0427, R2 = 0.138, F(1, 19) = 3.038, p = 0.0975)

## Discussion

Here we have created an alternative approach to SP prediction, hoping to take advantage of the new generation of computational tools to create quantitative *in silico* predictions of SP effectiveness. Such an approach could have a positive influence on various fields utilizing secreted proteins, including recombinant protein production and gene therapy. We chose to first test the approach with human BMP2, an important osteogenic protein long used in regenerative approach in bone both as a recombinant protein and a gene therapy. Notably hBMP2 is known to be a difficult-to-express protein [69, 70], and it was hoped SP engineering might help to alleviate this issue. One previous attempt at SP engineering in hBMP2 has been made using more traditional approaches but was unable to identify any SPs offering improved performance[18].

The first major choice in designing the new approach was how broad the computational screen would be and what assumptions would be made to choose the initial data set. There are no established strongly conserved motifs in eukaryotic SRP SPs[1], and as such there were few rules that could be applied when choosing the starting dataset. The eventual core assumptions employed were that only SRP SPs should be employed (i.e. no SPs from proteins that use alternative secretory pathways) and that SPs from closely related proteins would be more likely to conserve SP activity when applied in a heterologous context. Consequently the TGF-β superfamily, which contains BMP2, was chosen as the initial dataset.

The initial stages of the pipeline used the most up to date versions of two leading SP prediction tools (SignalP[60], DeepLoc[71]) to identify which SP-hBMP2 fusions were still likely to be secreted to the extracellular space. The results indicate these tools likely performed well, with all the tested signals being secreted to the extracellular space in both lines. However, current SP predictors are not capable of making quantitative predictions of how different SPs will perform in the same context. To try to address this problem a nucleotide sequence based approach was employed. As the signal peptide coding region contains the start codon and lies only slightly downstream of the ribosomal attachment point, secondary structure it forms can have a pronounced influence on translation. There is evidence that less stable secondary structure in a ±15bp window around the translational start site (which includes the beginning of the SP coding sequence) can improve translation initiation and early elongation[33–35]. Outside the translational start site mRNAs with a higher global stability appear to show improved translation, thought to be due to a combination of improved half-life and the generally higher stability of sequences containing many codons with abundant tRNAs[72, 73]. Consequently, it was though that heterologous SP-BMP2 fusions with mRNAs predicted to display less stable secondary structure at the translation start site and higher global stability might show improved translation and thus more protein export. An mRNA structure and stability based metric, SP score, was devised to assess these two properties simultaneously. It was hypothesised that SP score would correlate with levels of secreted protein.

To test the approach a small subset of the top results from the pipeline and two manually selected sequences were subjected to more detailed *in silico* analysis and tested *in vitro*. The top five candidates from the pipeline (termed SPs 1-5) were derived from avian, piscine and mammalian species, with SPs drawn from BMP15, GDF10 and three ARTN related genes (see table 1). The two manually selected sequences were a codon optimised variant of the native hBMP2 SP, and the human IL2 SP. A codon optimised hBMP2 SP allowed examination of the influence of nucleotide sequence changes divorced from changes to the protein sequence. In this case the tool TISIGNER was used to optimise the sequence for reduced opening energy[62], rather than a traditional codon adaption index (cAI) or tRNA adaption index (tAI) based approach. The hIL2 SP was chosen as it is a well-established choice in SP engineering, having been demonstrated to improve secretion for a variety of other proteins[74, 14, 75–77]. Nucleotide sequence alignment indicated there was modest homology between the top 5 TGF-β dataset candidates and the native hBMP2 SP, though the three ARTN orthologs contained a ∼25bp insertion. Amino acid sequence alignment showed that only a loose leucine rich motif appeared conserved through the set. Detailed analysis with SignalP 6 indicated that the insertion in the ARTN SPs was predicted to elongate the SP N-region, and that the loosely conserved leucine motif lay in the SP H-region, consistent with previous observations by others[78]. A final interesting prediction was that the hIL2 SP cleavage site would lie three residues downstream from the true SP/BMP2 proprotein boundary, though the prediction was made with lower confidence than in the other sequences. It was anticipated this could result in cleavage at this incorrect site with possible negative consequences for secretion and/or mature protein function, and possible cleavage site diversity.

*In vitro* testing in HEK293T showed SPs 1-5 showed slightly increased mean protein secretion versus the pVax_BMP2 positive control, however the data were highly variable and none were significantly higher than the control (see figure 4A). Of the two manually selected sequences hBMP2-TISIGNER showed almost no change versus pVax_BMP2 while hIL2 showed a modest and non-significant decrease. It was suspected that the HEK293T data variability was due to their tendency to form aggregates, leading to increased error during cell seeding which in turn influenced transfection unpredictably[79]. Despite none of the SPs inducing a significant increase in secretion, a correlation between SP score and secreted protein was found (see figure 5A). This was a promising result and suggested that the mRNA structure predictions had been accurate and had some influence on SP efficacy. Further work with a larger dataset would be required to confirm this fully.

C2C12 ELISA data showed that *in silico* sequences 1 & 2 and the manually selected BMP2 TISIGNER and hIL2 SPs induce significantly lower secretion BMP2 than the positive control, while SPs 3,4 & 5 showed non-significant decreases (see figure 4B). This was a clear change from the HEK293T results where there had been little difference between any of the sequences, and SP 2 had shown the highest mean concentration. The massive decrease in secretion seen in the two manually selected sequences was particularly interesting. In the case of SP hBMP2 TIS this indicate that nucleotide sequence was vitally important in this context, as a small number of synonymous mutations almost completely obviated secretion. SP hBMP2 TIS was predicted to show very similar opening energy to the native SP sequence, indicating that either the predictions had been inaccurate or additional factors were at play. Change in codon availability from HEK293T to C2C12 was considered, as the lines are human and murine respectively. A cAI based analysis was performed using the tool ATGme[80] and the murine codon usage table (http://www.kazusa.or.jp/codon/cgi-bin/showcodon.cgi?species=10090), indicating an addition of a single additional low availability codon (<10%) in the SP hBMP2 TIS versus the native sequence. This was thought to be unlikely to be the sole cause of the massive drop in secretion. An additional possible cause was interaction with antisense oligonucleotides or proteins that are found in C2C12 and not HEK293T. Further investigation would be required to identify these factors.

The SP hIL2 result was also notable as this marks a rare occurrence of the SP not offering improved performance versus a native SP, and indeed reducing it massively[74–77, 81]. It was suspected that this might be related to the predicted change in SP cleavage site for the SP hIL2 fusion. It is well established that multiple SP cleavage sites can be observed using one protein and cell type[30, 82, 83], therefore perhaps cleavage at both sites was seen in both lines but the balance between the two sites (or potentially even more than two sites) was shifted. Alternatively, cleavage may have occurred at the correct site but the SP introduced a more subtle alteration to early folding. Why this might occur in C2C12 and apparently not in HEK293T was unknown. Further work using LC-MS/MS would be able to empirically establish the cleavage site/s[30, 82, 83]. These two results further emphasise the importance of context in SP prediction and suggest that C2C12 may be somehow unusual in comparison to other lines commonly used in SP studies. Finally, no correlation was found between SP score and BMP2 secretion in C2C12 (see figure 5B).

ALP assay results at day 7 again showed background levels of ALP activity in all groups except the osteogenesis positive control (see figure 4C). This indicated that the novel SPs alone were not sufficient to induce an osteogenic effect in C2C12s. This observation was in line with previous data, where BMP2 is known to require either a high concentration or co-expression with other factors such as BMP7 to strongly induce osteogenesis[84, 66, 85, 86]. Combined with the ELISA data this was a clear indication that the SPs tested here were not sufficient to improve vector performance.

There were several clear weaknesses with the approach that were imposed by current limitations in computational methods. While 5’ end mRNA structure likely does play a role in how an SP influences protein secretion, it is certainly not the only factor. Interactions between the SP, SRP machinery and propeptide are known to be important [87–90], however making relevant predictions related to these interactions is difficult with existing computational tools and no established methods exist. While recent advances in multimer tertiary structure prediction by tools such as AlphaFold2 are impressive and have seen some use in modelling SP/translocon interactions[91, 92], modelling of the kinetics of the interactions of multiple multi-subunit complexes is not within current capabilities. If tools able to produce reliable predictions of these interactions are developed, they could perhaps be combined with mRNA structure predictions to improve efficacy. The initial limitation of the sequence panel to the TGF-β superfamily dataset was an attempt to address this problem. It was hoped that SPs from these proteins would be more likely to retain their function in a new but similar context, though it is possible that this was not a reasonable assumption and the dataset was too diverse.

It should also be noted that while RNA structure prediction has seen continuous improvement and a similar burst in progress with machine learning based approaches, predictions are still frequently inaccurate in many contexts[40]. This is thought to be due to biases in the annotated RNA structure data sets used for model training, which are frequently dominated by short ncRNAs and are unable to account for all possible biomacromolecule interactions[93]. Consequently, the mRNA structure predictions used in this approach must be taken with a reasonable degree of scepticism. Advances in high through RNA structure determination techniques promise to address these problems[94, 95, 40], and the ever growing volume of training data from wider contexts promises to allow continued improvement of RNA structure modelling in future. Despite this, the significant correlation observed between SP score and secreted protein in the HEK293T data suggests the predictions were somewhat accurate, though further work would be required to conclusively establish this was not due to chance. Future work could employ a library of synonymous variants to separate the influence of nucleotide sequence from protein structure.

## Conclusions

The mRNA structure-based method for SP effectiveness prediction described here was capable of identifying previously untested SPs capable of functioning comparably to the native sequence in HEK293T. Additionally there was a significant correlation between model predictions and secreted protein in HEK293T, though none of the SPs showed a significant improvement in secretion versus the native SP. The approach was not effective in C2C12 cells with several SPs inducing a significant decrease in secretion. Particularly poor performance from the codon optimised SP hBMP2 TIS indicated the importance of synonymous mutations in the SP and merits further study. These results suggest the mRNA structure prediction approach requires further improvement before it can produce significant improvements in this context, and ideally should be combined more precise protein sequence based predictions of SP activity in future.

## List of abbreviations

4NP: 4-Nitrophenol
ALP: Alkaline Phosphetase
BMP2: Bone Morphogenetic Protein 2 MFE Minimum Free Energy
NPP: Nitrophenol Phosphate
rhBMP2: Recombinant human Bone Morphogenetic Protein 2
Sec/SPI: Sec substrates cleaved by SPase I
SP: Signal Peptide
SRP: Signal Recognition Particle
TISIGNER: Translation initiation coding region designer

## Declarations

### Ethics approval and consent to participate

Not applicable

### Consent for publication

Not applicable

### Availability of data and materials

The datasets used and analysed during the study were publicly available and access links are provided in the main text. Full results from the *in silico* screen can be provided on request.

### Authors’ contributions

PW conceived the study, planned, and performed the computational screen, planned and performed the *in vitro* validation experiments, analysed the data, wrote the manuscript and prepared the manuscript figures.

BJ provided advice during study planning and with molecular biology, provided the HEK293T cells, and reviewed the manuscript.

HF provided access to lab space and equipment for the *in vitro* work, and reviewed the manuscript.

DW, PG & RD oversaw the experiments and reviewed the manuscript.

## Other acknowledgements

The authors would like to acknowledge Professors David Wood (Oral Biology, Faculty of Medicine and Health, University of Leeds) and Peter Giannoudis (Academic Department of Trauma and Orthopaedics, School of Medicine, University of Leeds) for their support during this work.

## Bibliography

1. Owji H, Nezafat N, Negahdaripour M, Hajiebrahimi A, Ghasemi Y. A comprehensive review of signal peptides: Structure, roles, and applications. European Journal of Cell Biology. 2018;97:422–41.

2. Meazza R, Gaggero A, Neglia F, Basso S, Sforzini S, Pereno R, et al. Expression of two interleukin-15 mRNA isoforms in human tumors does not correlate with secretion: role of different signal peptides. European Journal of Immunology. 1997;27:1049–54.

3. Schröder M. Engineering eukaryotic protein factories. Biotechnology Letters. 2008;30:187–96.

4. Stern B, Olsen LC, Tröße C, Ravneberg H, Pryme IF. Improving mammalian cell factories: The selection of signal peptide has a major impact on recombinant protein synthesis and secretion in mammalian cells. Trends Cell Mol Biol. 2007;2:1–17.

5. Kober L, Zehe C, Bode J. Optimized signal peptides for the development of high expressing CHO cell lines. Biotechnology and Bioengineering. 2013;110:1164–73.

6. Delic M, Göngrich R, Mattanovich D, Gasser B. Engineering of Protein Folding and Secretion—Strategies to Overcome Bottlenecks for Efficient Production of Recombinant Proteins. Antioxidants & Redox Signaling. 2014;21:414–37.

7. Zahrl RJ, Prielhofer R, Ata Ö, Baumann K, Mattanovich D, Gasser B. Pushing and pulling proteins into the yeast secretory pathway enhances recombinant protein secretion. Metabolic Engineering. 2022;74:36–48.

8. You M, Yang Y, Zhong C, Chen F, Wang X, Jia T, et al. Efficient mAb production in CHO cells with optimized signal peptide, codon, and UTR. Applied Microbiology and Biotechnology. 2018;102:5953–64.

9. Ohmuro-Matsuyama Y, Yamaji H. Modifications of a signal sequence for antibody secretion from insect cells. Cytotechnology. 2018;70:891–8.

10. Zou Z, Wang R, Go EP, Desaire H, Sun PD. High level stable expression of recombinant HIV gp120 in glutamine synthetase gene deficient HEK293T cells. Protein Expression and Purification. 2021;181:105837–105837.

11. Wilkinson C, Kyle J, Irimpen M, Stuart S, Mohandass S, Sheperd A, et al. Improved yield of recombinant human IFN-α2b from mammalian cells using heterologous signal peptide approach. Protein Expression and Purification. 2022;198:106125–106125.

12. Mullin MJ, Wilkinson C, Hiles I, Smith KJ. Improved secretion of recombinant human IL-25 in HEK293 cells using a signal peptide-pro-peptide domain derived from Trypsin-1. Biotechnology Letters. 2021. 10.1007/s10529-020-03072-z.

13. Román R, Miret J, Scalia F, Casablancas A, Lecina M, Cairó JJ. Enhancing heterologous protein expression and secretion in HEK293 cells by means of combination of CMV promoter and IFNα2 signal peptide. Journal of Biotechnology. 2016;239:57–60.

14. Zhang L, Leng Q, Mixson AJ. Alteration in the IL-2 signal peptide affects secretion of proteins in vitro and in vivo. The Journal of Gene Medicine. 2005;7:354–65.

15. Sun B, Zhang H, Benjamin Jr. DK, Brown T, Bird A, Young SP, et al. Enhanced Efficacy of an AAV Vector Encoding Chimeric, Highly Secreted Acid α-Glucosidase in Glycogen Storage Disease Type II. Molecular Therapy. 2006;14:822–30.

16. Ma H, Liu Y, Liu S, Xu R, Zheng D. Oral adeno-associated virus-sTRAIL gene therapy suppresses human hepatocellular carcinoma growth in mice. Hepatology. 2005;42:1355–63.

17. Roberts SA, Dong B, Firrman JA, Cao W, Xiao W. 462. Engineering the Human Factor VIII Signal Peptide for Enhanced Secretion. Molecular Therapy. 2014;22:S177–S177.

18. Hacobian AR, Posa-Markaryan K, Sperger S, Stainer M, Hercher D, Feichtinger GA, et al. Improved osteogenic vector for non-viral gene therapy. European cells & materials. 2016;31:191–204.

19. Egan KP, Awasthi S, Tebaldi G, Hook LM, Naughton AM, Fowler BT, et al. A Trivalent HSV-2 gC2, gD2, gE2 Nucleoside-Modified mRNA-LNP Vaccine Provides Outstanding Protection in Mice against Genital and Non-Genital HSV-1 Infection, Comparable to the Same Antigens Derived from HSV-1. Viruses. 2023;15.

20. Loomis RJ, DiPiazza AT, Falcone S, Ruckwardt TJ, Morabito KM, Abiona OM, et al. Chimeric Fusion (F) and Attachment (G) Glycoprotein Antigen Delivery by mRNA as a Candidate Nipah Vaccine. Front Immunol. 2021;12:772864.

21. Billerhart M, Schönhofer M, Schueffl H, Polzer W, Pichler J, Decker S, et al. CD47-targeted cancer immunogene therapy: Secreted SIRPα-Fc fusion protein eradicates tumors by macrophage and NK cell activation. Molecular Therapy - Oncolytics. 2021;23:192–204.

22. Tsuchiya Y, Morioka K, Taneda I, Shirai J, Yoshida K. Gene design of signal sequence for the effective secretion of recombinant protein using insect cell. Nucleic Acids Symposium Series. 2005;49:305–6.

23. Knappskog S, Ravneberg H, Gjerdrum C, Tröβe C, Stern B, Pryme IF. The level of synthesis and secretion of Gaussia princeps luciferase in transfected CHO cells is heavily dependent on the choice of signal peptide. Journal of Biotechnology. 2007;128:705–15.

24. Cheng K-W, Wang F, Lopez GA, Singamsetty S, Wood J, Dickson PI, et al. Evaluation of artificial signal peptides for secretion of two lysosomal enzymes in CHO cells. Biochem J. 2021;478:2309–19.

25. Bamford RN, DeFilippis AP, Azimi N, Kurys G, Waldmann TA. The 5′ Untranslated Region, Signal Peptide, and the Coding Sequence of the Carboxyl Terminus of IL-15 Participate in Its Multifaceted Translational Control1. The Journal of Immunology. 1998;160:4418–26.

26. Figueiredo Neto M, Figueiredo ML. Skeletal muscle signal peptide optimization for enhancing propeptide or cytokine secretion. Journal of Theoretical Biology. 2016;409:11–7.

27. Haryadi R, Ho S, Kok YJ, Pu HX, Zheng L, Pereira NA, et al. Optimization of Heavy Chain and Light Chain Signal Peptides for High Level Expression of Therapeutic Antibodies in CHO Cells. PLOS ONE. 2015;10:e0116878–e0116878.

28. Srila W, Baumann M, Borth N, Yamabhai M. Codon and signal peptide optimization for therapeutic antibody production from Chinese hamster ovary (CHO) cell. Biochemical and Biophysical Research Communications. 2022;622:157–62.

29. Peng C, Guo Y, Ren S, Li C, Liu F, Lu F. SPSED: A Signal Peptide Secretion Efficiency Database. Frontiers in Bioengineering and Biotechnology. 2022;9.

30. Park J-H, Lee H-M, Jin E-J, Lee E-J, Kang Y-J, Kim S, et al. Development of an in vitro screening system for synthetic signal peptide in mammalian cell-based protein production. Appl Microbiol Biotechnol. 2022;106:3571–82.

31. Holec PV, Camacho KV, Breuckman KC, Mou J, Birnbaum ME. Proteome-Scale Screening to Identify High-Expression Signal Peptides with Minimal N-Terminus Biases via Yeast Display. ACS Synth Biol. 2022;11:2405–16.

32. Ding Y, Tang Y, Kwok CK, Zhang Y, Bevilacqua PC, Assmann SM. In vivo genome-wide profiling of RNA secondary structure reveals novel regulatory features. Nature. 2014;505:696–700.

33. Meredith C, Amanda S, Gabriela P, Lela L, Benjamin Z, A. VH, et al. An RNA structure-mediated, posttranscriptional model of human α-1-antitrypsin expression. Proceedings of the National Academy of Sciences. 2017;114:E10244–53.

34. Mustoe AM, Busan S, Rice GM, Hajdin CE, Peterson BK, Ruda VM, et al. Pervasive Regulatory Functions of mRNA Structure Revealed by High-Resolution SHAPE Probing. Cell. 2018;173:181–195.e18.

35. Mustoe AM, Corley M, Laederach A, Weeks KM. Messenger RNA Structure Regulates Translation Initiation: A Mechanism Exploited from Bacteria to Humans. Biochemistry. 2018;57:3537–9.

36. Xiang Y, Huang W, Tan L, Chen T, He Y, Irving PS, et al. Pervasive downstream RNA hairpins dynamically dictate start-codon selection. Nature. 2023;621:423–30.

37. Wang Y, Mao Y, Xu X, Tao S, Chen H. Codon Usage in Signal Sequences Affects Protein Expression and Secretion Using Baculovirus/Insect Cell Expression System. PLOS ONE. 2015;10:e0145887–e0145887.

38. Cheng Y, Liu S, Lu C, Wu Q, Li S, Fu H, et al. Missense mutations in the signal peptide of the porcine GH gene affect cellular synthesis and secretion. Pituitary. 2016;19:362–9.

39. Xu Y, Verma D, Sheridan RP, Liaw A, Ma J, Marshall NM, et al. Deep Dive into Machine Learning Models for Protein Engineering. J Chem Inf Model. 2020;60:2773–90.

40. Zhang J, Fei Y, Sun L, Zhang QC. Advances and opportunities in RNA structure experimental determination and computational modeling. Nature Methods. 2022;19:1193– 207.

41. Teufel F, Almagro Armenteros JJ, Johansen AR, Gíslason MH, Pihl SI, Tsirigos KD, et al. SignalP 6.0 predicts all five types of signal peptides using protein language models. Nature Biotechnology. 2022;40:1023–5.

42. Almagro Armenteros JJ, Sønderby CK, Sønderby SK, Nielsen H, Winther O. DeepLoc: prediction of protein subcellular localization using deep learning. Bioinformatics. 2017;33:3387–95.

43. Sato K, Akiyama M, Sakakibara Y. RNA secondary structure prediction using deep learning with thermodynamic integration. Nature Communications. 2021;12:941.

44. Katagiri T, Watabe T. Bone Morphogenetic Proteins. Cold Spring Harb Perspect Biol. 2016;8.

45. Bostrom MPG. Expression of Bone Morphogenetic Proteins in Fracture Healing. Clinical Orthopaedics and Related Research®. 1998;355.

46. Huntley R, Jensen E, Gopalakrishnan R, Mansky KC. Bone morphogenetic proteins: Their role in regulating osteoclast differentiation. Bone Reports. 2019;10:100207.

47. Sánchez-Duffhues G, Hiepen C, Knaus P, ten Dijke P. Bone morphogenetic protein signaling in bone homeostasis. Bone. 2015;80:43–59.

48. Wang RN, Green J, Wang Z, Deng Y, Qiao M, Peabody M, et al. Bone Morphogenetic Protein (BMP) signaling in development and human diseases. Genes & Diseases. 2014;1:87– 105.

49. Marsell R, Einhorn TA. The role of endogenous bone morphogenetic proteins in normal skeletal repair. Injury. 2009;40 Suppl 3:S4–7.

50. Khan SN, Lane JM. The use of recombinant human bone morphogenetic protein-2 (rhBMP-2) in orthopaedic applications. Expert Opinion on Biological Therapy. 2004;4:741– 8.

51. Schlundt C, Bucher CH, Tsitsilonis S, Schell H, Duda GN, Schmidt-Bleek K. Clinical and Research Approaches to Treat Non-union Fracture. Current osteoporosis reports. 2018;16:155–68.

52. Carragee EJ, Hurwitz EL, Weiner BK. A critical review of recombinant human bone morphogenetic protein-2 trials in spinal surgery: emerging safety concerns and lessons learned. The Spine Journal. 2011;11:471–91.

53. James AW, LaChaud G, Shen J, Asatrian G, Nguyen V, Zhang X, et al. A Review of the Clinical Side Effects of Bone Morphogenetic Protein-2. Tissue Eng Part B Rev. 2016;22:284–97.

54. Wilkinson P, Bozo IY, Braxton T, Just P, Jones E, Deev RV, et al. Systematic Review of the Preclinical Technology Readiness of Orthopedic Gene Therapy and Outlook for Clinical Translation. Frontiers in Bioengineering and Biotechnology. 2021;9:39–39.

55. UniProt: the universal protein knowledgebase in 2021. Nucleic Acids Research. 2021;49:D480–9.

56. Wheeler DL, Barrett T, Benson DA, Bryant SH, Canese K, Chetvernin V, et al. Database resources of the National Center for Biotechnology Information. Nucleic Acids Research. 2007;35 suppl_1:D5–12.

57. Cunningham F, Allen JE, Allen J, Alvarez-Jarreta J, Amode MR, Armean IM, et al. Ensembl 2022. Nucleic Acids Research. 2022;50:D988–95.

58. Cummins C, Ahamed A, Aslam R, Burgin J, Devraj R, Edbali O, et al. The European Nucleotide Archive in 2021. Nucleic Acids Research. 2022;50:D106–10.

59. Davis P, Zarowiecki M, Arnaboldi V, Becerra A, Cain S, Chan J, et al. WormBase in 2022—data, processes, and tools for analyzing Caenorhabditis elegans. Genetics. 2022;220:iyac003.

60. Teufel F, Almagro Armenteros JJ, Johansen AR, Gíslason MH, Pihl SI, Tsirigos KD, et al. SignalP 6.0 predicts all five types of signal peptides using protein language models. Nature Biotechnology. 2022;40:1023–5.

61. Almagro Armenteros JJ, Sønderby CK, Sønderby SK, Nielsen H, Winther O. DeepLoc: prediction of protein subcellular localization using deep learning. Bioinformatics. 2017;33:3387–95.

62. Bhandari BK, Lim CS, Gardner PP. TISIGNER.com: web services for improving recombinant protein production. Nucleic Acids Research. 2021;49:W654–61.

63. Sato K, Akiyama M, Sakakibara Y. RNA secondary structure prediction using deep learning with thermodynamic integration. Nature Communications. 2021;12:941.

64. Lorenz R, Bernhart SH, Höner zu Siederdissen C, Tafer H, Flamm C, Stadler PF, et al. ViennaRNA Package 2.0. Algorithms for Molecular Biology. 2011;6:26–26.

65. Kerpedjiev P, Hammer S, Hofacker IL. Forna (force-directed RNA): Simple and effective online RNA secondary structure diagrams. Bioinformatics. 2015;31:3377–9.

66. Feichtinger GA, Hacobian A, Hofmann AT, Wassermann K, Zimmermann A, van Griensven M, et al. Constitutive and inducible co-expression systems for non-viral osteoinductive gene therapy. European cells & materials. 2014;27:166–84; discussion 184.

67. Zhao B, Katagiri T, Toyoda H, Takada T, Yanai T, Fukuda T, et al. Heparin potentiates the in vivo ectopic bone formation induced by bone morphogenetic protein-2. Journal of Biological Chemistry. 2006;281:23246–53.

68. Kim MG, Kim CL, Kim YS, Jang JW, Lee GM. Selective endocytosis of recombinant human BMPs through cell surface heparan sulfate proteoglycans in CHO cells: BMP-2 and BMP-7. Scientific Reports. 2021;11:3378.

69. Jérôme V, Thoring L, Salzig D, Kubick S, Freitag R. Comparison of cell-based versus cell-free mammalian systems for the production of a recombinant human bone morphogenic growth factor. Engineering in Life Sciences. 2017;17:1097–107.

70. Riedl SAB, Jérôme V, Freitag R. Repeated Transient Transfection: An Alternative for the Recombinant Production of Difficult-to-Express Proteins Like BMP2. Processes. 2022;10.

71. Thumuluri V, Almagro Armenteros JJ, Johansen AR, Nielsen H, Winther O. DeepLoc 2.0: multi-label subcellular localization prediction using protein language models. Nucleic Acids Research. 2022;50:W228–34.

72. Presnyak V, Alhusaini N, Chen Y-H, Martin S, Morris N, Kline N, et al. Codon optimality is a major determinant of mRNA stability. Cell. 2015;160:1111–24.

73. Mauger DM, Cabral BJ, Presnyak V, Su SV, Reid DW, Goodman B, et al. mRNA structure regulates protein expression through changes in functional half-life. Proceedings of the National Academy of Sciences. 2019;116:24075 LP – 24083.

74. Bamford RN, DeFilippis AP, Azimi N, Kurys G, Waldmann TA. The 5′ Untranslated Region, Signal Peptide, and the Coding Sequence of the Carboxyl Terminus of IL-15 Participate in Its Multifaceted Translational Control1. The Journal of Immunology. 1998;160:4418–26.

75. Loomis RJ, DiPiazza AT, Falcone S, Ruckwardt TJ, Morabito KM, Abiona OM, et al. Chimeric Fusion (F) and Attachment (G) Glycoprotein Antigen Delivery by mRNA as a Candidate Nipah Vaccine. Frontiers in Immunology. 2021;12.

76. Billerhart M, Schönhofer M, Schueffl H, Polzer W, Pichler J, Decker S, et al. CD47-targeted cancer immunogene therapy: Secreted SIRPα-Fc fusion protein eradicates tumors by macrophage and NK cell activation. Mol Ther Oncolytics. 2021;23:192–204.

77. Egan KP, Awasthi S, Tebaldi G, Hook LM, Naughton AM, Fowler BT, et al. A Trivalent HSV-2 gC2, gD2, gE2 Nucleoside-Modified mRNA-LNP Vaccine Provides Outstanding Protection in Mice against Genital and Non-Genital HSV-1 Infection, Comparable to the Same Antigens Derived from HSV-1. Viruses. 2023;15.

78. Nilsson I, Lara P, Hessa T, Johnson AE, von Heijne G, Karamyshev AL. The Code for Directing Proteins for Translocation across ER Membrane: SRP Cotranslationally Recognizes Specific Features of a Signal Sequence. Journal of Molecular Biology. 2015;427:1191–201.

79. Sticky Issues with 293 Cells. https://www.culturecollections.org.uk/news/ecacc-news/sticky-issues-with-293-cells.aspx. Accessed 25 Oct 2023.

80. Daniel E, Onwukwe GU, Wierenga RK, Quaggin SE, Vainio SJ, Krause M. ATGme: Open-source web application for rare codon identification and custom DNA sequence optimization. BMC Bioinformatics. 2015;16:303.

81. Zhang L, Leng Q, Mixson AJ. Alteration in the IL-2 signal peptide affects secretion of proteins in vitro and in vivo. The Journal of Gene Medicine. 2005;7:354–65.

82. Huang Y, Fu J, Ludwig R, Tao L, Bongers J, Ma L, et al. Identification and quantification of signal peptide variants in an IgG1 monoclonal antibody produced in mammalian cell lines. J Chromatogr B Analyt Technol Biomed Life Sci. 2017;1068–1069:193–200.

83. Haryadi R, Ho S, Kok YJ, Pu HX, Zheng L, Pereira NA, et al. Optimization of Heavy Chain and Light Chain Signal Peptides for High Level Expression of Therapeutic Antibodies in CHO Cells. PLOS ONE. 2015;10:e0116878–e0116878.

84. Kawai M, Bessho K, Maruyama H, Miyazaki J, Yamamoto T. Simultaneous gene transfer of bone morphogenetic protein (BMP) -2 and BMP-7 by in vivo electroporation induces rapid bone formation and BMP-4 expression. BMC Musculoskelet Disord. 2006;7:62.

85. Feichtinger GA, Hofmann AT, Slezak P, Schuetzenberger S, Kaipel M, Schwartz E, et al. Sonoporation increases therapeutic efficacy of inducible and constitutive BMP2/7 in vivo gene delivery. Human gene therapy methods. 2014;25:57–71.

86. Kaito T, Morimoto T, Mori Y, Kanayama S, Makino T, Takenaka S, et al. BMP-2/7 heterodimer strongly induces bone regeneration in the absence of increased soft tissue inflammation. Spine J. 2018;18:139–46.

87. Bornemann T, Jöckel J, Rodnina MV, Wintermeyer W. Signal sequence-independent membrane targeting of ribosomes containing short nascent peptides within the exit tunnel. Nat Struct Mol Biol. 2008;15:494–9.

88. Zhang X, Schaffitzel C, Ban N, Shan S. Multiple conformational switches in a GTPase complex control co-translational protein targeting. Proceedings of the National Academy of Sciences. 2009;106:1754–9.

89. Jungnickel B, Rapoport TA. A posttargeting signal sequence recognition event in the endoplasmic reticulum membrane. Cell. 1995;82:261–70.

90. Zhang X, Rashid R, Wang K, Shan S. Sequential Checkpoints Govern Substrate Selection During Cotranslational Protein Targeting. Science. 2010;328:757–60.

91. Bryant P, Pozzati G, Elofsson A. Improved prediction of protein-protein interactions using AlphaFold2. Nature Communications. 2022;13:1265.

92. Gutierrez Guarnizo SA, Kellogg MK, Miller SC, Tikhonova EB, Karamysheva ZN, Karamyshev AL. Pathogenic signal peptide variants in the human genome. NAR Genom Bioinform. 2023;5:lqad093.

93. Danaee P, Rouches M, Wiley M, Deng D, Huang L, Hendrix D. bpRNA: large-scale automated annotation and analysis of RNA secondary structure. Nucleic Acids Res. 2018;46:5381–94.

94. Siegfried NA, Busan S, Rice GM, Nelson JAE, Weeks KM. RNA motif discovery by SHAPE and mutational profiling (SHAPE-MaP). Nature Methods. 2014;11:959–65.

95. Smola MJ, Weeks KM. In-cell RNA structure probing with SHAPE-MaP. Nature Protocols. 2018;13:1181–95.

